# Prediction of fruit texture with training population optimization for efficient genomic selection in apple

**DOI:** 10.1101/862193

**Authors:** Morgane Roth, Mario Di Guardo, Walter Guerra, Hélène Muranty, Andrea Patocchi, Fabrizio Costa

**Affiliations:** Plant Breeding Research Division, Agroscope, Wädenswil, Switzerland; Department of Genomics and Biology of Fruit Crops, Research and Innovation Centre, Fondazione Edmund Mach (FEM), Via E. Mach 1, 38010 San Michele all’Adige, Italy; Dipartimento di Agricoltura, Alimentazione e Ambiente, University of Catania, Catania, Italy; Research Centre Laimburg, Laimburg 6, 39040 Auer, Italy; IRHS, INRAE, Agrocampus-Ouest, Université d’Angers, SFR 4207 QuaSaV, Beaucouzé, France

**Author notes:** Corresponding author: Morgane Roth. Further e-mails: Mario Di Guardo; Walter Guerra; Hélène Muranty; Andrea Patocchi; Fabrizio Costa.

**Keywords:** Apple, genomic prediction, rrBLUP, multi-trait, fruit texture, relatedness, training set optimization

## Abstract

Texture plays a major role in the determination of fruit quality in apple. Due to its physiological and economic relevance, this trait has been largely investigated, leading to the fixation of the major gene PG1 controlling firmness in elite cultivars. To further improve fruit texture, the targeting of an undisclosed reservoir of loci with minor effects is compelling. In this work, we aimed to unlock this potential with a genomic selection approach by predicting fruit acoustic and mechanical features as obtained with a TA.XT*plus* texture analyzer in 537 individuals genotyped with 8,294 SNP markers. The best prediction accuracies following cross-validations within the training set (TRS) of 259 individuals were obtained for the acoustic linear distance (0.64). Prediction accuracy was further improved through the optimization of TRS size and composition according to the test set. With this strategy, a maximal accuracy of 0.81 was obtained when predicting the synthetic trait PC1 in the family ‘Gala × Pink Lady’. We discuss the impact of genetic relatedness and clustering on trait variability and predictability. Moreover, we demonstrated the need for a comprehensive dissection of the complex texture phenotype and the potentiality of using genomic selection to improve fruit quality in apple.

**Highlight:** A genomic selection study, together with the optimization of the training set, demonstrated the possibility to accurately predict texture sub-traits valuable for the amelioration of fruit quality in apple.

## Introduction

Fruits, during maturation and ripening, undergo a complex series of genetically programmed events contributing to their attractiveness and suitability for human consumption. Amongst the various physiological and physical changes, fruit texture is certainly the most important and investigated traits, especially in apple. A favorable texture is in fact highly appreciated by consumers, enabling, moreover, a long-term storage.

Texture can nowadays be dissected into two groups of sub-traits, mechanical and acoustic, contributing to distinguish between firm (based on mechanical sub-traits) and crispy (based on acoustic sub-traits) types of apples. These texture parameters have been already described and validated in apple (Costa *et al*., 2011, 2012), and were implemented in QTL-mapping studies carried out with bi-parental populations (Longhi *et al*., 2012) as well as more structured approaches, such as Pedigreed Based Analysis (PBA) and Genome-Wide Association Studies (GWAS, Kumar *et al*., 2013; Migicovsky *et al*., 2016; Amyotte *et al*., 2017; Di Guardo *et al*., 2017; McClure *et al.*, 2019). These works elucidated the complex genetic control of the fruit texture in apple, identifying a large number of QTLs distributed over the apple genome, with the most relevant regions located on chromosome 3, 10 and 16. This genetic complexity is moreover reflected in the regulation of the cell-wall and middle lamella disassembling, a physiological process orchestrated by a myriad of cell-wall modifying enzymes (Giovannoni, 2001; Costa *et al*., 2010a). This highly polygenic control can hamper the selection assisted by molecular markers in breeding activities programmed to ameliorate fruit texture performance (Iwata *et al*. 2016). In the QTL mapping studies carried out to date, a major region was located on chromosome 10, close to the polygalacturonase locus (Costa *et al.*, 2010b; Longhi *et al.*, 2013). This QTL explains a high (about 40%) yet incomplete part of the texture variance, leaving room for better harnessing this trait. As introduced by Di Guardo *et al*. (2017), in modern breeding programs this locus has been fixed through successive rounds of *ad-hoc* crossing and selection. In turn, the phenotypic variance of modern families, obtained by crossing valuable parents for texture performance, might now be under the control of other loci with minor-effect. Selection based on QTLs associated to this trait can therefore be limited by the fact that QTL-based approaches ignore small effect QTLs possibly underlying the control of such traits (Desta & Ortiz, 2014). To face this limitation, an alternative approach for genome-assisted breeding known as genomic selection (GS) has been introduced by the seminal work of Meuwissen *et al.* (2001). In contrast to marker assisted selection, GS defines the estimation of the genetic merit of an individual taking into account all genome-wide distributed genetic markers, making it especially relevant for complex traits (Heffner *et al.*, 2009). GS considers two sets of individuals: the training set (TRS), genotyped and phenotyped to train a prediction model, and the test set (TS, also called validation set), represented by individuals only genotyped on which the genomic estimated breeding value (GEBVs) is estimated (Heffner *et al.*, 2009; Crossa *et al.*, 2017). In principle, the most favorable scenario for GS is to predict highly heritable traits in a TS highly related to the TRS. While trait heritability can be increased (to a certain extent) by more precise and more repeated phenotyping, relatedness between TS and TRS can be optimized with different strategies. Dedicated approaches and tools have been proposed to address this issue based on optimization parameters (Laloë, 1993; Rincent *et al.*, 2012; Isidro *et al.*, 2015) and algorithms (Akdemir *et al.*, 2015). In theory, it could thus be feasible to acquire phenotypic and genotypic data for a highly diverse TRS in the first place and, in the second place, to retain individuals of the optimal TRS for a given TS *in silico*.

GS has been largely applied in major crops for primary traits such as yield (Crossa *et al.*, 2017). In perennial species, GS would have a great potential in improving the breeding efficiency due to their long generation time (McClure *et al.*, 2014). It has been pioneered in forest trees (reviewed in Grattapaglia, 2017) and more recently in fruit trees such as crops from the *Malus, Citrus and Pyrus* genera (Muranty *et al.* 2015; Minamikawa *et al.*, 2017, 2018). GS has also been recently employed to investigate fruit quality in tomato (Duangjit *et al.*, 2016), while in apple standard fruit pomological traits were predicted using 8 to 20 full-sib families as training populations (Kumar *et al.*, 2012, 2015; Muranty *et al.*, 2015). In apple, low to high prediction accuracies were obtained depending on the cross-validation design and on trait heritability. Among these studies, fruit texture was only partially addressed via classical fruit firmness measurements (Kumar *et al.*, 2012, 2015; McClure *et al.*, 2018) and sensorial evaluation (Kumar *et al.*, 2015).

In this work, we attempted to predict fruit texture in 6 full-sib families with a diverse training set considering several acoustic and mechanical traits dissecting fruit texture. Further, we explored the methodological improvements that can be made to optimize the TRS according to the TS, which contributed to improve prediction accuracies. In this context, we discussed the feasibility of genomic selection for ameliorating fruit quality through molecular assisted breeding programs.

## Materials and methods

### Plant Material

The plant material and phenotyping strategies used in this work have been detailed in previous works (Costa *et al.*, 2011; Longhi *et al.*, 2012; Di Guardo *et al.*, 2017). Briefly, two types of plant materials have been used in this survey. The first was an apple collection represented by 259 accessions planted in three replicates at the experimental orchards of the Fondazione Edmund Mach (Trento) in the Northern part of Italy. The second type of plant material consisted of 6 full-sib biparental families, for a total of 278 offsprings. Two (‘FjDe’: ‘Fuji’ x ‘Delearly’ and ‘FjPL’: ‘Fuji’ x ‘Pink Lady’) were located at the Fondazione Edmund Mach (same orchard as the collection), while the other four (‘GaPL’: ‘Royal Gala’ x ‘Pink Lady’, ‘GaPi’: ‘Royal Gala’ x ‘Pinova’, ‘FjPi’: ‘Fuji’ x ‘Pinova’ and ‘GDFj’: ‘Golden Delicious’ x ‘Fuji’) were planted at the experimental orchard of the Laimburg Research Center (Bolzano), located in the same area with near-identical climatic and pedological conditions. At the time of the analysis, all plants (from both collection and families, together named as ‘population’ here) were in a productive and adult phase. Fruit texture was phenotyped in 2012, 2013 and 2015 for the collection, and in 2012 and 2013 for ‘FjDe’ and ‘FjPL’ and in 2012 and 2014 for the four remaining families (Table 1). Unlike the collection, each offspring belonging to the six families was represented by a single tree (no replicates). All plants, from both collection and bi-parental families, were grafted on ‘M9’ rootstock and grown according to conventional horticultural management for plant training, pruning and pest-disease control.

**Table 1.** Description of the whole population and experimental design used for genomic prediction of texture. Maternal and paternal cultivars are given for full-sib biparental families. FEM, Foundation Edmund Mach; RCL, Research Center Laimburg. Cluster assignments as given by the discriminant analysis of principal components on 8,294 markers. Relationship to collection, mean additive relationship of progenies relative to collection.

| Name | Mother | Father | Location | Evaluated years | # IDs | Cluster assignments<br>(# IDs) |  |  |  |  |  | Relationship to collection |
| --- | --- | --- | --- | --- | --- | --- | --- | --- | --- | --- | --- | --- |
|  |  |  |  |  |  | 1 | 2 | 3 | 4 | 5 | 6 |  |
| FjDe | Fuji | Delearly | FEM | 2012-13 | 50 | 8 | 20 | 0 | 0 | 22 | 0 | -0.056 |
| FjPi | Fuji | Pinova | RCL | 2012-14 | 70 | 1 | 30 | 0 | 0 | 39 | 0 | -0.078 |
| FjPL | Fuji | Pink Lady | FEM | 2012-13 | 80 | 0 | 50 | 0 | 0 | 30 | 0 | -0.071 |
| GaPi | Royal Gala | Pinova | RCL | 2012-14 | 36 | 0 | 0 | 0 | 0 | 36 | 0 | -0.021 |
| GaPL | Royal Gala | Pink Lady | RCL | 2012-14 | 15 | 0 | 0 | 0 | 0 | 15 | 0 | -0.020 |
| GDFj | Golden Delicious | Fuji | RCL | 2012-14 | 27 | 0 | 6 | 0 | 0 | 21 | 0 | -0.057 |
| Collection | - | - | FEM | 2012-13-15 | 259 | 45 | 37 | 31 | 55 | 66 | 25 | - |

Fruits were harvested from each plant at the time of the physiological ripening stage, established according to standard horticultural fruit quality parameters, such as the change in color of the skin, seeds and flesh, fruit firmness value and the iodine coloration index indicating the internal starch degradation. After harvest, fruits were stored for two months at 2°C with 95% of relative humidity.

### Texture phenotyping

The texture performance of the apple fruit was phenotypically dissected into mechanical and acoustic sub-traits with the use of a texture analyzer TA.XT*plus* (Stable MicroSystems Ltd., Godalming, UK) equipped with an acoustic envelop device AED (Stable MicroSystems Ltd., Godalming, UK), as described in Costa *et al.* (2011). For each genotype included in the population, a homogeneous set of five apples was collected. Four identical discs were isolated per fruit, avoiding seeds, seed cavity tissues or skin, for a total of 20 measurements per genotype (5 biological replicates and 4 technological replicates). Each texture profile was then digitally elaborated identifying 12 texture measurements (*i. e.* ‘sub-traits’), four related to the acoustic performance and eight to the mechanical force-displacement. In brief, the mechanical sub-traits were coded as: initial, final, maximum and mean force (related to the different force values associated to the different parts of the force-displacement profile), area, force linear distance (derived length of the profile), Young’s module (also known as elasticity module) and number of force peaks. The acoustic sub-traits were maximum and mean acoustic pressure, acoustic linear distance and number of acoustic peaks. A more exhaustive and complete description of the texture sub-traits is reported in Costa *et al.* 2011.

### SNP genotyping

The DNA employed for the genotyping of each individual considered in this survey was isolated from young leaves collected at the beginning of the vegetative phase with the Qiagen DNeasy Plant Kit and further quantified with a Nanodrop ND-8000 (ThermoScientific, USA). SNP markers were genotyped through the HiScan (Illumina, USA) and the apple 20K SNP chip Infinium array (Illumina, USA) assembled within the framework of the European project FruitBreedomics (Bianco *et al.*, 2014). The SNP pattern was initially analyzed with the software GenomeStudio and further re-edited with ASSiST (Di Guardo *et al.*, 2015). SNPs with minor allele frequencies lower than 0.05 and call rate below 0.2 were filtered out with the package ‘snpStats’ (Clayton, 2019). The final set of markers successfully recovered in the population consisted in 8,294 biallelic SNPs.

### Analysis of the fruit texture sub-traits

We used a mixed linear model to get the best linear unbiased predictors (BLUPs) of each individual’s genotypic value. For each apple measured, we first calculated the mean over the four technical replicates to retain only the biological replication level in the model. Each of the twelve mechanical or acoustic sub-traits, considered as ‘Y’, was explained by the genotype as random effect, the trial (location by year) as fixed effect and the random effect of the error as: *Y*_*i,j,k*_ = *µ* + *genotype*_*i*_ + *trial*_*j*_ + *e*_*i,j,k*_ (1), with each phenotypic datapoint *Y*_*i,j,k*_ explained by the mean *µ*, the genotype i, the trial j and the error for each combination of genotype, trial and replicate (k, *i.e.* a single apple). This model was fitted separately for all traits with the ‘lme4’ R-package (Bates *et al.*, 2015). Broad-sense heritability was calculated as 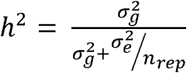 (2), where 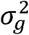 is the genotypic variance, 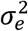 is the error variance and *n*_*rep*_ the mean number of repetitions.

Principal component analysis (PCA) was performed on BLUPs with the ‘FactorMiner’ R-package (Lê *et al.*, 2008). Only values from the collection were used to create the principal components, while the families were plotted as supplementary individuals with principal components (PC) coordinates calculated on the base of the PCs initially built with the collection. Coordinates of individuals on the first and the second PCs (‘PC1’ and ‘PC2’) were used for prediction and subsequently named ‘synthetic’ traits.

### Kinship and clustering analyses

The realized additive relationship was calculated with the ‘A.mat’ function of the ‘rrBLUP’ package (Endelman, 2011) and depicted in a heatmap plot obtained with the R-function ‘heatmap.2’ (package ‘gplots’, Warnes *et al.*, 2016). Genetic clustering was further assessed in the collection with a discriminant analysis of principal components (DAPC, Jombart *et al.*, 2010), carried out with the R-package ‘adegenet’ (Jombart, 2008) using the entire set of 8,294 markers. In the first step, six significant clusters were retained with the function ‘find.clusters’ using 300 principal components and selecting the number of clusters with the highest likelihood (based on the Bayesian information criterion value-BIC, Fig. S1). Out of these variables, 150 were retained and employed in the clustering computed with the ‘dapc’ function, which created five principal components that maximized the inter-cluster distance while minimizing the inter-individual distance within each cluster. The assignment of offsprings to clusters was obtained with the function ‘predict_dapc’. Pairwise Fst values between clusters were then computed with the entire SNP set with the function ‘pairwise. WCfst’ from R-package ‘hierfstat’ (Yang, 1998, Goudet 2005).

### Prediction models

Genomic predictions were computed through two models implemented in the rrBLUP framework, as reported in Endelman *et al.* 2011 (and ‘rrBLUP R’-package):

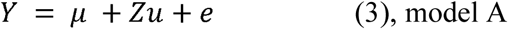

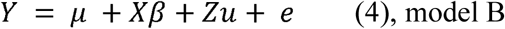

where Y is the vector of BLUPs of the genotypic values (*n* × 1), *µ* is the mean of the phenotype, W is the *n* × *p* incidence matrix linking the genotypes to observations of Y, G contains the allelic states of the marker loci (additive coding −1,0,1), *u* the *p* × 1 vector of random marker effects with 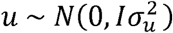, and e is a *n* × 1 vector of random errors. Model (B), contains also X, the *n* × c incidence matrix for cluster assignment of each individual, where c is the number of clusters and *β* is the *c* × 1 vector of the cluster fixed effects.

A 5-fold cross-validation was applied within the collection with both model (A) and (B) respectively and repeated 100 times. For predicting each family (considered then as TS), three different TRS composition rules, named as “scenarios”, were tested using the two models *without a priori genetic information on individuals*. In scenario 1, each family was predicted using the collection only. In scenario 2, 30% of individuals of the predicted family were instead added to the collection in the TRS while the remaining 70% formed the TS. In scenario 3, a single half-sib family (*e.g*. ‘GaPL’ is half-sib with ‘FjPL’ and ‘GaPi’) was added to the collection to form the TRS, leading to two to four TRS possibilities (and accuracy values). To illustrate the scenarios taking ‘GaPi’ as an example, scenario 1 corresponded to [TRS = COLL // TS = ‘GaPi’] (one accuracy estimation only), scenario 2 corresponded to [TRS = 30% ‘GaPi’ offsprings + COLL // TS = 70 % remaining offsprings of ‘GaPi’] (sampling of the 30% repeated 100 times, giving 100 estimations of the accuracy), and scenario 3 corresponded to [TRS = ‘GaPL’ *or* ‘FjPi’ + COLL // TS = ‘GaPi’] (resulting here in the estimation of two accuracy values).

TRS optimization was then performed *with a priori genetic information on individuals* by varying TRS size with different optimization methods relying on the prediction model A. To this end, a relatedness-driven and a principal component-driven approaches were adopted. The relatedness-driven approach was tested in three different manners: (i) by starting with the 10 most-related individuals and adding single individuals with decreasing mean relationship to the family, or (ii) with decreasing maximum relationship to the family (N=10 to N=259), or (iii) by starting with a TRS composed of the most related cluster and adding less and less related clusters successively (final TRS size N=259). In the principal component-driven approach, TRS individuals were selected with increasing TRS size using a protocol by Akdemir (R-package ‘STPGA’, 2019). The optimal TRS with increasing size from 10 individuals to 259 with increments of 20 individuals was chosen based on the five principal components obtained with DAPC analysis and using the ‘CDmean’ design criteria and the function ‘GenAlgForSubsetSelection’. Here, individuals were chosen independently for each TRS size, meaning that we did not proceed to a gradual enrichment of the TRS.

All accuracy values were based on Pearson correlation calculated between observed values (*i.e.* BLUPs of genotypic values) and predicted values of the TS individuals. When standard deviations were not available, we calculated an approximate 95% confidence interval of the correlation coefficient with a Fisher’s Z-transformation (‘cor.test’ function in base R). Calculations were performed in R (R Core Team, 2014) and graphs were created with the R-package ‘ggplot2’ (Wickham, 2016).

## Results

### Fruit texture phenotypic dissection

The fruit texture phenotypic data used in this survey were represented by the analysis of multi-trait features accurately dissected into 4 acoustic and 8 mechanical sub-traits (Table 2, Table S1). A mixed linear model was used to obtain BLUPs of genotypic values used in the further analyses. The texture sub-traits showed an overall high heritability, spanning from 0.90-0.96 for the entire population (collection and families) to 0.88-0.94 for the apple accessions included in the collection (Table 2). In order to visualize the diversity and inheritance of fruit texture profiles, a principal component analysis (PCA) was performed using the twelve textural sub-traits measured in the collection, while individuals from families were considered as supplementary individuals (see also Di Guardo et al. 2017, Fig. 1). In this analysis, the first PC axis (PC1), explaining 80.5% of phenotypic variability, comprehensively summarizing the general variability of the twelve phenotypic variables. The second axis (PC2), instead, mainly differentiated the acoustic from mechanical sub-traits, explaining a smaller, yet substantial, portion of the phenotypic variability (12.7%, Fig. 1A).

**Table 2.** Summary of texture traits assessed in the whole population. h^2^, broad sense heritability. For comparison, h^2^ are also given considering measurements of the collection only.

| Trait | Mean | SD | $h^2$ | $h^2$<br>(COLL only) |
| --- | --- | --- | --- | --- |
| ALD | 5094 | 2049 | 0.938 | 0.928 |
| ANP | 50.4 | 39.1 | 0.924 | 0.909 |
| APMax | 65.2 | 4.38 | 0.921 | 0.880 |
| APMean | 49.6 | 3.12 | 0.951 | 0.915 |
| Area | 813 | 273 | 0.951 | 0.930 |
| FF | 10.1 | 3.98 | 0.946 | 0.929 |
| FLD | 101 | 5.78 | 0.955 | 0.937 |
| Fmax | 11.8 | 4.02 | 0.946 | 0.924 |
| FMean | 9.60 | 3.31 | 0.951 | 0.929 |
| FNP | 17.9 | 4.16 | 0.934 | 0.931 |
| IF | 9.94 | 3.29 | 0.933 | 0.906 |
| YM | 1.19 | 0.353 | 0.899 | 0.890 |

**Figure 1.**
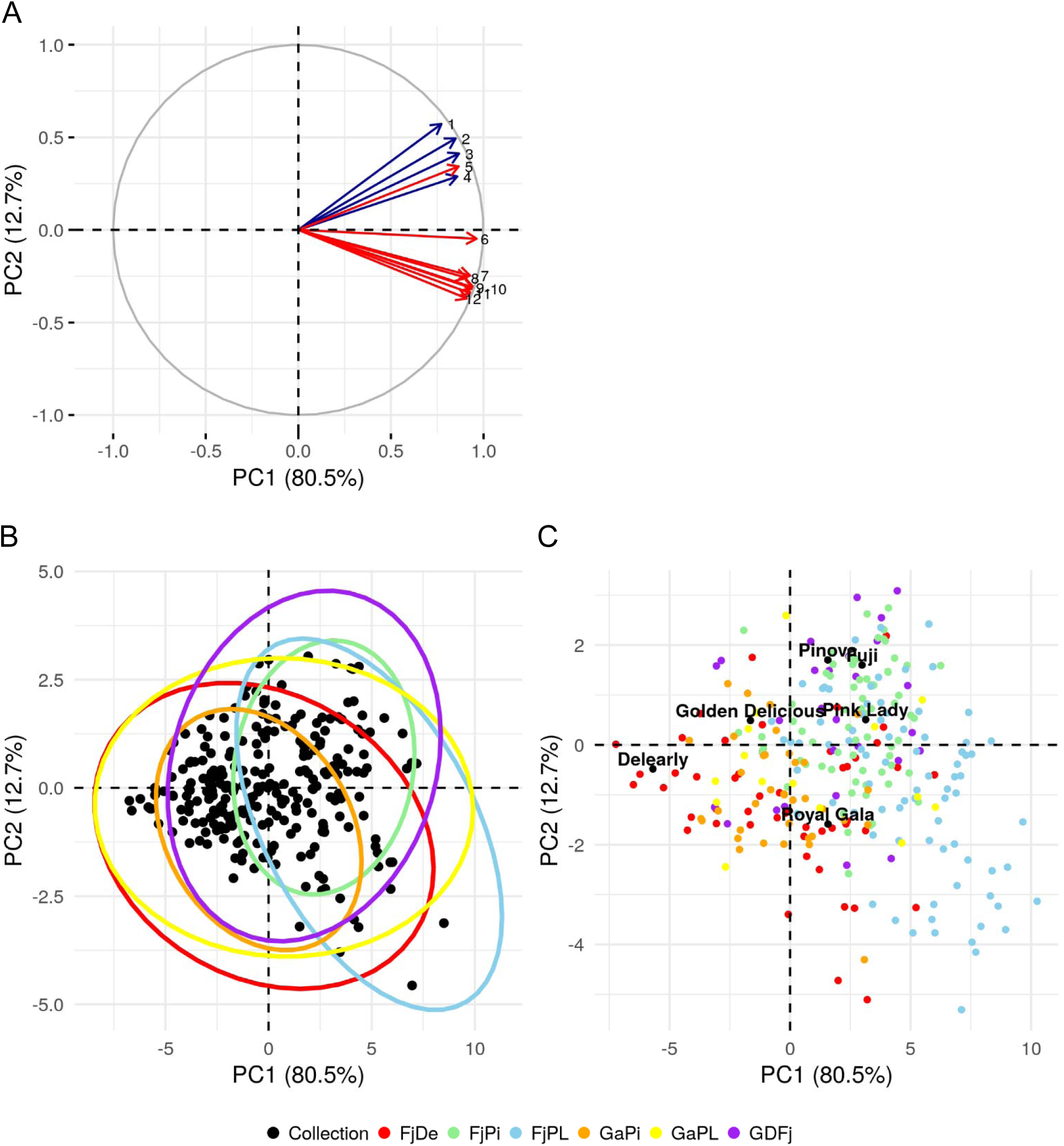
Principal component analysis (PCA) of 12 texture sub-traits. A, PCA 2D-plot of variables, with acoustic traits in blue and mechanical traits in red with 1, ANP; 2, ALD; 3, APMAx; 4, APMean; 5, FNP; 6, FLD; 7,FF; 8, YM; 9, Area; 10; Fmean; 11, Fmax; 12, IF. B, PCA 2D-plot of individuals with collection individuals represented as dots and families as ellipses. C, PCA 2D-plot of individuals showing family offspring and their respective parents.

In the distinction between the two types of texture sub-traits (mechanical and acoustic) by PC2, it is worth noting that one mechanical variable (FNP) was oriented together with the acoustic group. FNP was in fact more correlated with acoustic sub-traits (mean correlation 0.77) than with the rest of the mechanical ones (mean correlation 0.69, Fig. 1A). Individuals of the population were present in the four quadrants of the PCA 2D-plot, identifying different types of texture: mealy (negative PC1), predominantly firm (positive PC1 and negative PC2) and predominantly crispy (positive PC1 and positive PC2, Fig. 1B). With this regard, the distribution of texture profiles indicated that the collection is mainly composed of individuals with low to moderate crispiness and firmness at the exception of few outliers. It is also important to note that variation on the PC2 axis is much lower for accessions having a negative PC1 value, illustrating that mealy apples cannot be crispy (Fig. 1B).

The six parental cultivars, known to have different texture profiles after two months of storage, were, as expected, plotted over the different quadrants of the PCA 2D-plot (Fig. 1C). ‘Delearly’ and ‘Golden Delicious’ were plotted in the area corresponding to the mealy type of apple, while ‘Royal Gala’ was instead grouped with moderately firm apples. ‘Fuji’, ‘Pink Lady’ and ‘Pinova’ were instead positioned in the positive quadrant for both PC1 and PC2, corresponding to the crispy type of apple. The populations originated by the controlled cross of these varieties were also distributed over the PCA plot with specific orientations (Fig. 1B**-** C). In particular, ‘FjPL’ offsprings were mostly projected towards the ‘firm quadrant’, while ‘GDFj’ was more oriented in the ‘crispy quadrant’ (Fig. 1B). Moreover, the segregation of the families was very variable with regard to their corresponding parental profiles (Fig. 1C). While ‘GDFj’ was the only family showing a classic type of segregation (intermediate between the parents), the distributions of the other families were more similar to one of the two parents (‘FjDe’ and ‘GaPi’), with a varying number of offsprings being of transgressive type (‘FjDe’, ‘GaPL’, ‘FjPi’ and ‘FjPL’). In particular, while ‘Fuji’ and ‘Pink Lady’ showed a very similar texture profile on PC1 (2.99 and 3.14 respectively), major differences were observed on the PC2 (1.6 and 0.51 respectively, Fig. 1C, Table S1). Variation in the texture performance of ‘FjPL’ offsprings was also observed on the PC2 axis, although with a much broader variation with regards to ‘Fuji’ and ‘Pink Lady’. Accordingly, apples of this family were overall firm to very firm while having a very low to very high crispiness (Fig. 1C, Table S1, Fig. S2).

### Additive relationship and genetic clustering in the population

The accuracy of genomic prediction is highly correlated to the level of relatedness between the training and the test sets (TRS and TS). To identify the overall patterns of relatedness between families and the collection, a clustering analysis of all the individuals based on their pairwise additive relationship was performed (Fig. 2). The parental cultivar ‘Royal Gala’ was found to be the most related to the rest of the collection (mean additive relatedness −6.32E-4), while ‘Fuji’ was the most distantly related (mean additive relatedness −0.102, Table S2). Accordingly, ‘Royal Gala’–related families were more closely related to the collection respect to the four ‘Fuji’-related families, plotted together on the top-right panel of the heatmap (Fig. 2). Mean additive relationship values for each family reflected the patterns observed on the heatmap, namely higher values for ‘GaPi’ and ‘GaPL’ (−0.021 to −0.020) and lower for ‘Fuji’-related families (−0.056 to −0.078, Table 1, Table S2).

**Figure 2:**
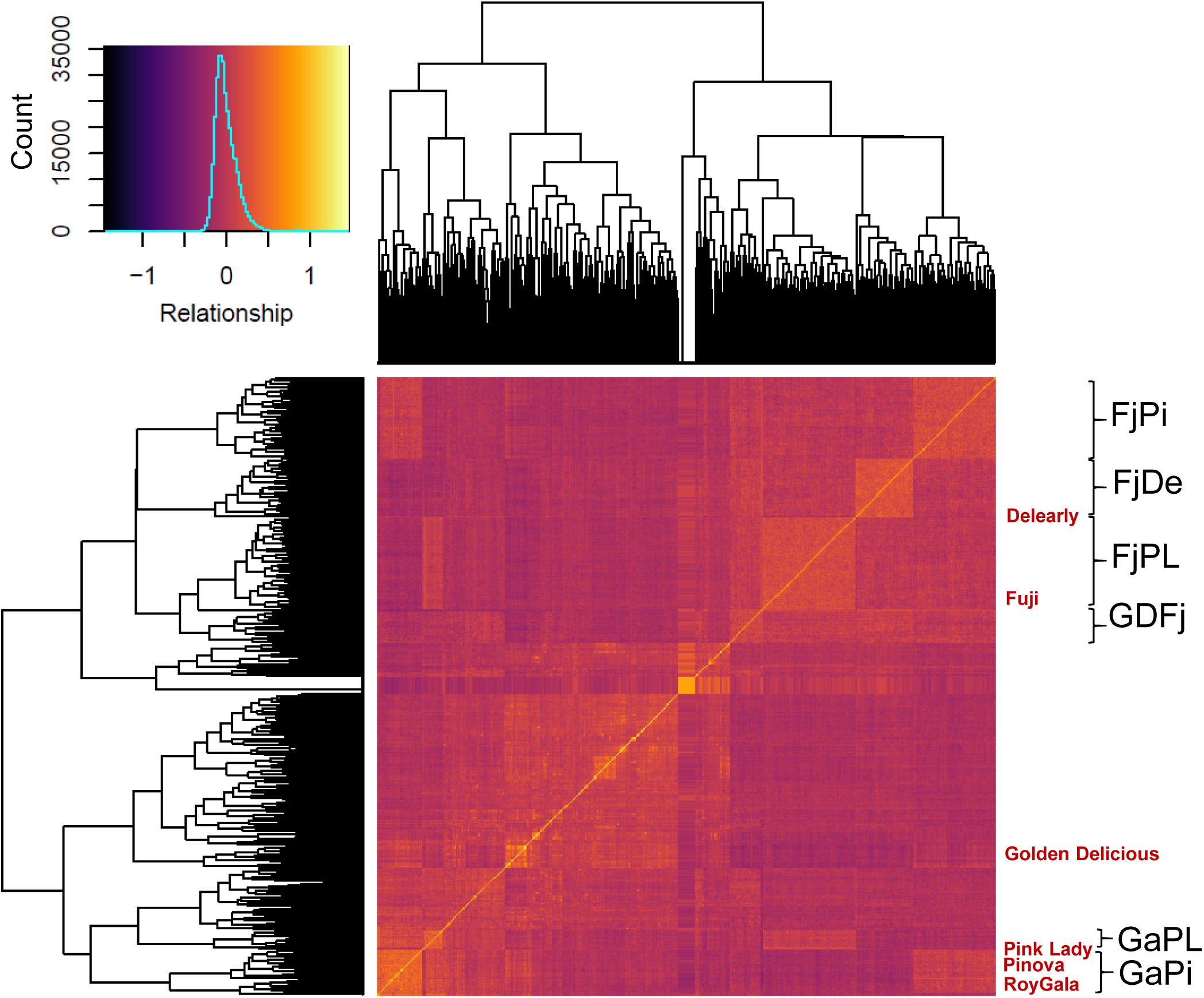
Realized additive relationship calculated with 8,294 SNPs. Families indicated in black with brackets and parents are indicated in red.

To investigate the genetic structure of the collection and its impact on the prediction accuracy, a discriminant analysis of principal component with the entire SNP set (8,294 SNPs) was performed. Through the BIC criteria, six clusters, described with five principal components, were defined as the most probable (see Methods, Fig. S1). All parental cultivars were assigned to cluster 5, except ‘Fuji’ that was grouped in cluster 2 (Fig. 3A, Table S3). Of these clusters, cluster 5 resulted to be the largest (N=66), while the smallest was cluster 6 (n=25, Table 1, Table S3). The cluster assignment in families was predicted using the principal components derived by the DAPC analysis carried out on the collection. Most of the individuals were assigned to the parental clusters 2 and 5, while 8 individuals of ‘FjDe’ and one of ‘FjPi’ were assigned to cluster 1 (Table 1, Fig. 3B-C). Overall, clusters 2 and 5 contained the largest part of the whole population, while clusters 1, 3, 4 and 6 were the lowest represented (Fig. 3C, Table S3). However, while the DAPC analysis suggested this genetic clustering as the most realistic in the diversity panel represented by the collection, the pairwise Fst-values between clusters indicated a low genetic differentiation (values comprised between 0.002 and 0.018, Table S4). The Fst value between clusters 2 and 5, containing the parents and most of their offsprings, was for instance 0.013. As our design allowed the comparison of families obtained from crosses within cluster 5 (‘Royal Gala’-related) and between clusters 2 and 5 (‘Fuji’-related), the information on genetic clustering was further used to control the genetic background in the subsequent prediction models (‘model B’, see Methods). The phenotypic distributions across clusters reveal that clusters 2 and 5 have, for all traits except PC2, elevated values compared to other clusters, with values of cluster 2 individuals surpassing those of cluster 5 (Fig. S3), indicating a possible correlation existing between genetic clustering and texture.

**Figure 3:**
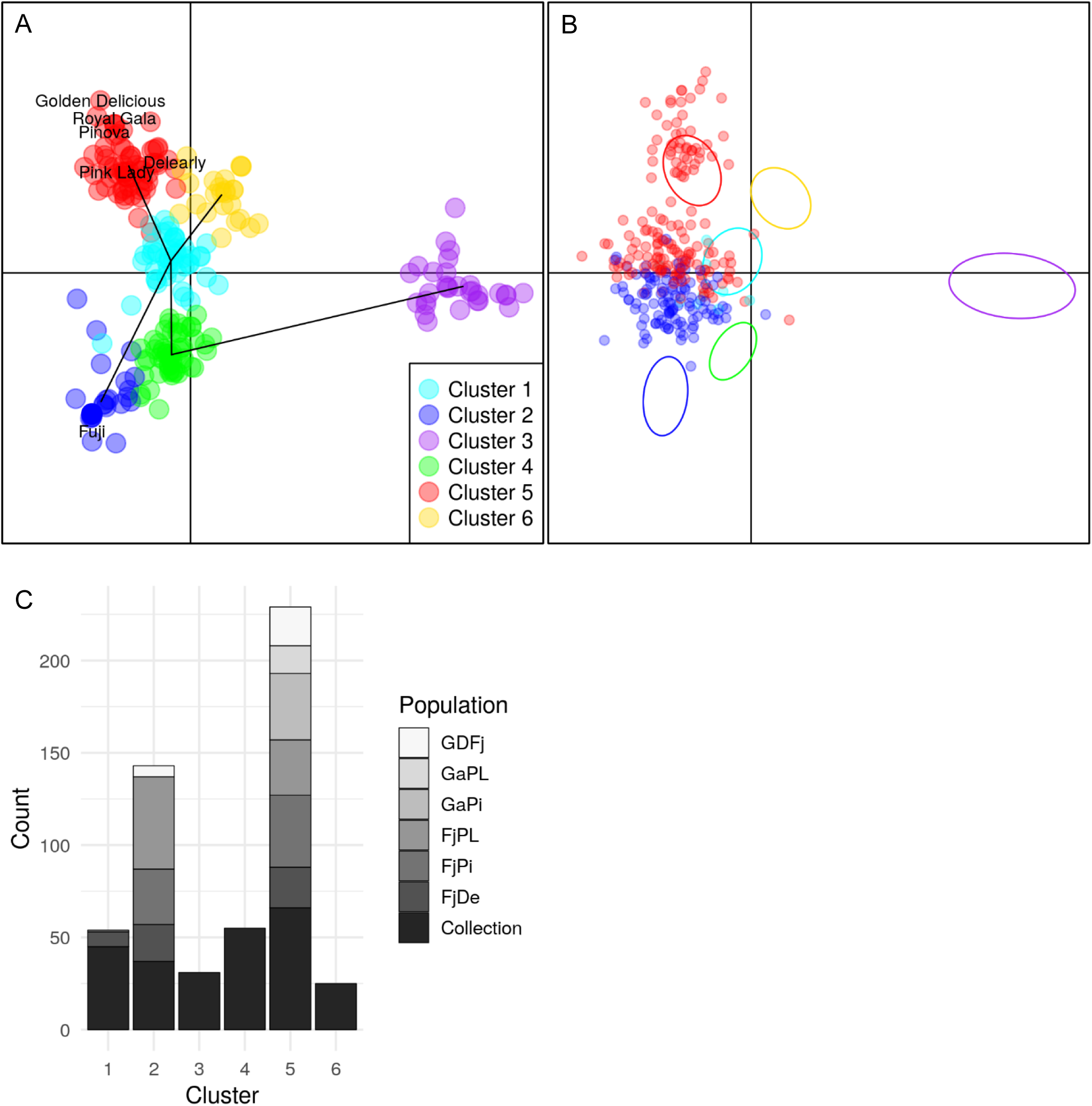
Discriminant analysis of principal components and cluster assignments of individuals based on 8,294 SNPs. A, projection on principal component (PC) 1 and 3 of the cluster assignments of individuals in the collection with parents of families indicated with their names. Black lines materialize the PCs defining clusters. B, Predicted cluster assignments of progenies of the six full-sib families projected on PC1 and PC3 axes and represented by dots, with collection individuals in the six genetic clusters represented as ellipses (same color legend as in part A). C, Distribution of individuals across the six genetic clusters in each population.

### Cross-validations within collection

A hundred 5-fold cross-validations within the collection were run with the additive rrBLUP model on BLUPs with and without considering the genetic clustering as a fixed effect (models A and B, respectively). In this context, PC1 and PC2 were also considered as traits, leading in the end to 14 predicted traits (Fig. S4). Instead of improving predictions, the inclusion of the clustering effect degraded accuracies for all traits, with a maximum accuracy decrease of 0.02 for the mean force (FMean). The highest mean prediction was obtained for the acoustic linear distance (ALD, *mean cor* = 0.64, Fig. S4) whereas the number of force peaks yielded the second highest accuracy (FNP, *mean cor* = 0.63, Fig. S4, Table S5). Moreover, while FNP yielded a relatively high accuracy as inferred from heritability (0.93, Table 2), the overall mean accuracies among traits did not follow the ranking of heritability obtained within the collection phenotypes (Wilcoxon signed-rank-test, p-value = 4.88E-4, model A).

### Genomic prediction of families without training population optimization

In practice, families can be predicted with any available related genetic material that has been genotyped and phenotyped. For this reason, three different scenarios of training population design were tested, including or not individuals from the predicted family or from a half-sib family (see Methods, “Prediction models”). The predictions in each of these scenarios were calculated with the two prediction models (A and B, respectively depicted in Fig. 4, Fig. S5). Without clustering, overall three families (‘FjPi’, ‘GaPi’ and ‘GaPL’) could be predicted with moderate to high accuracies (accuracies ranging from 0.08 for PC2 in ‘GaPi’ to 0.73 for PC1 in ‘GaPL’, respectively), with PC1 being the best predicted trait among these families (mean for scenario 1, model A: 0.50, Fig. 4). The three remaining families yielded near-zero (‘FjPL’) or negative accuracies (‘FjDe’ and ‘GDFj’, mean accuracies between −0.29 and 0.30, Fig. 4). The correlations between predicted and observed values for each individual and for all traits and families obtained are depicted in Fig. S6 (model A and scenario 1). Out of 252 combinations of trait, scenario and family predictions, only 74 gave better accuracies (considering an increase in accuracy larger than 0.01). When considering accuracies above 0.20, this number dropped to 40 out of 103 family-trait-scenario combinations (maximum gain: 0.04, Table S6, Fig 4, Fig. S5). Thus, the implementation of clustering did not clearly improve the predictions of families.

**Figure 4:**
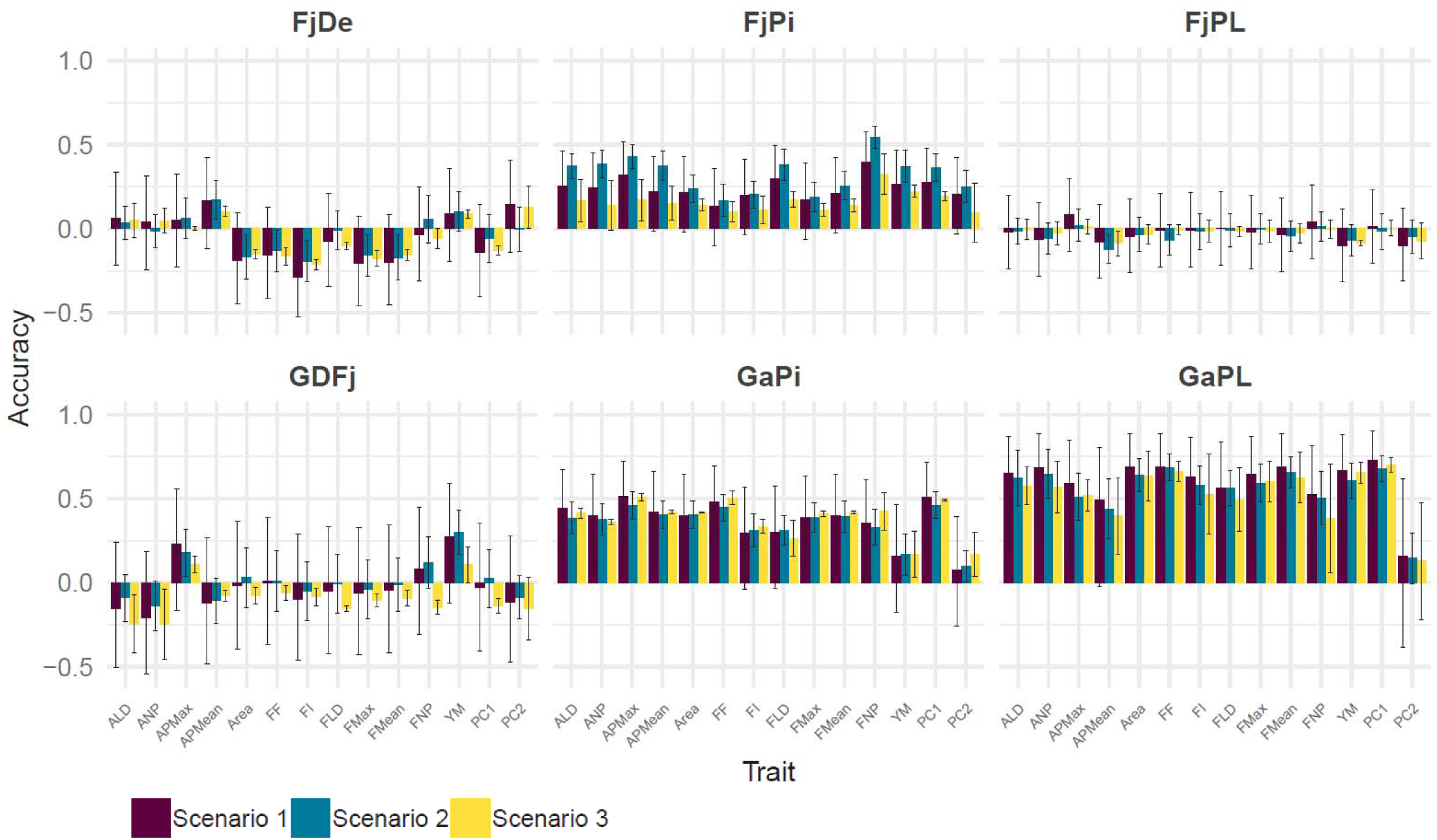
Mean and standard deviation of accuracies obtained in three prediction scenarios. In scenario 1, each family was predicted using the collection only. In scenario 2, 30% of individuals of the predicted family were added to the collection in the TRS and the remaining 70% formed the TS. In scenario 3, a single half-sib family was added to the collection to form the TRS. The predictions were made with model A, which does not take into account the genetic clustering of individuals.

It is also important to underline that the addition of related individuals to the collection did not systematically improve the predictions. For instance, in ‘GaPL’ the prediction was more accurate with scenario 1 with regards to scenario 2 and 3 (mean prediction accuracies of 0.60, 0.56 and 0.53 respectively for scenario 1, 2, 3, respectively, model A). Scenario 2 particularly improved the accuracies in ‘FjPi’ (mean accuracies of 0.32, model A) as it better predicted 12 out of 14 traits. Scenario 3 instead was the lowest performing, although it increased the prediction accuracy of 7 traits (8 with clustering) in ‘GaPi’ (mean accuracy of 0.38, all values across trait in model A, Fig 4, Table S6).

### Genomic prediction of families with training population optimization

To test the hypothesis that retaining only the most related individuals or clusters in the TRS might allow to maximize prediction accuracies, we compared the predictive abilities obtained for each family and trait using training sets with different sizes. This process started with a small TRS having the highest relatedness to which individuals were added in the order of decreasing relatedness to reach the size of the entire collection using three different enrichment procedures (see Methods). TRS optimization was also carried out with a more sophisticated approach based on the optimization algorithm presented by Akdemir *et al.* (Akdemir *et al.*, 2015; Akdemir & Isidro-Sánchez, 2019), using DAPC-defined principal components and the ‘CD-Mean’ value as decision criterion. The results obtained using these different methods are illustrated in Table 3, Fig. 5 and Fig. S7 for four traits selected for their practical relevance (ALD, FNP, PC1 and PC2) while results for the remaining traits are reported in Table S7. Regarding the four selected traits, the best accuracy for each of the 6 × 4 family-trait combinations was in most cases obtained with the addition of single individuals based on their relationship to the family (in 10 cases using the maximum relationship and in 10 cases using the mean relationship, Fig. 5A and B, Table 3, Table S7). The mean optimal population size was 92 individuals with a minimum size of 10 and a maximum size of 202 individuals (Table 3, Table S7), meaning that the entire collection was never considered as the optimal TRS for predicting texture. The maximal accuracies observed ranged from 0.01 to 0.81, which corresponded to a mean increase in accuracy of 0.17 when compared to predictions of families with the entire collection (minimum increase: 0.02; maximum increase: 0.40 – compared to scenario 1, model A). The highest accuracy was 0.81, and was obtained for the “multi-trait” PC1 in ‘GaPL’ family with only 129 individuals, *i.e.* nearly half of the collection size. The distribution of accuracies with increasing TRS size in each family for the four focal traits was also investigated (Fig. 5). Overall, traits tended to follow the same trend within a family. In families ‘GaPL’ and ‘GaPi’, which had the highest relatedness to the collection among all families (Table 1), the accuracy was moderate to high from as few as 100 individuals for ALD, FNP and PC1, and remained relatively stable while increasing TRS size (Fig. 5A-D). ‘FjPi’ was the only family for which increasing TRS up to 200 individuals resulted in a clear accuracy improvement, with any of the approaches implemented here (Fig. 5A-D). In families with overall low accuracies, such as ‘FjDe’, ‘FjPL’ and ‘GDFj’, the highest accuracy was in most cases obtained with 10 to 70 individuals, and declined or remained stable with larger TRS size (Fig. 5A-D). In ‘GDFj’, for instance, accuracies above 0.2 were found only with a TRS of 10 to 66 individuals (Fig. 5A-C, Table S7). Moreover, while FNP was not predictable in ‘GDFj’ with the entire collection (*cor* = 0.08 for *N* = 259), an improved accuracy of 0.32 was observed with as few as 15 individuals (based on maximum relationship, Fig. 5B).

**Table 3.** Maximum accuracies obtained among four training set optimization methods in predictions made for each combination of trait and family.

| Family | Trait | Accuracy | TRS size | Method |
| --- | --- | --- | --- | --- |
| FjDe | ALD | 0.23 | 77 | Mean relationship |
| FjDe | FNP | 0.18 | 21 | Max relationship |
| FjDe | PC1 | 0.26 | 77 | Mean relationship |
| FjDe | PC2 | 0.36 | 56 | Max relationship |
| FjPi | ALD | 0.36 | 189 | Mean relationship |
| FjPi | FNP | 0.59 | 174 | Max relationship |
| FjPi | PC1 | 0.36 | 178 | Max relationship |
| FjPi | PC2 | 0.26 | 202 | Mean relationship |
| FjPL | ALD | 0.10 | 130 | CDmean-opt |
| FjPL | FNP | 0.16 | 22 | Max relationship |
| FjPL | PC1 | 0.20 | 120 | Max relationship |
| FjPL | PC2 | 0.22 | 10 | CDmean-opt |
| GaPi | ALD | 0.46 | 156 | Mean relationship |
| GaPi | FNP | 0.40 | 13 | Max relationship |
| GaPi | PC1 | 0.54 | 191 | Clusters |
| GaPi | PC2 | 0.28 | 19 | Mean relationship |
| GaPL | ALD | 0.72 | 136 | Clusters |
| GaPL | FNP | 0.78 | 37 | Max relationship |
| GaPL | PC1 | 0.81 | 129 | Mean relationship |
| GaPL | PC2 | 0.40 | 140 | Max relationship |
| GDFj | ALD | 0.21 | 66 | Mean relationship |
| GDFj | FNP | 0.32 | 15 | Max relationship |
| GDFj | PC1 | 0.19 | 31 | Mean relationship |
| GDFj | PC2 | 0.01 | 10 | Mean relationship |

**Figure 5:**
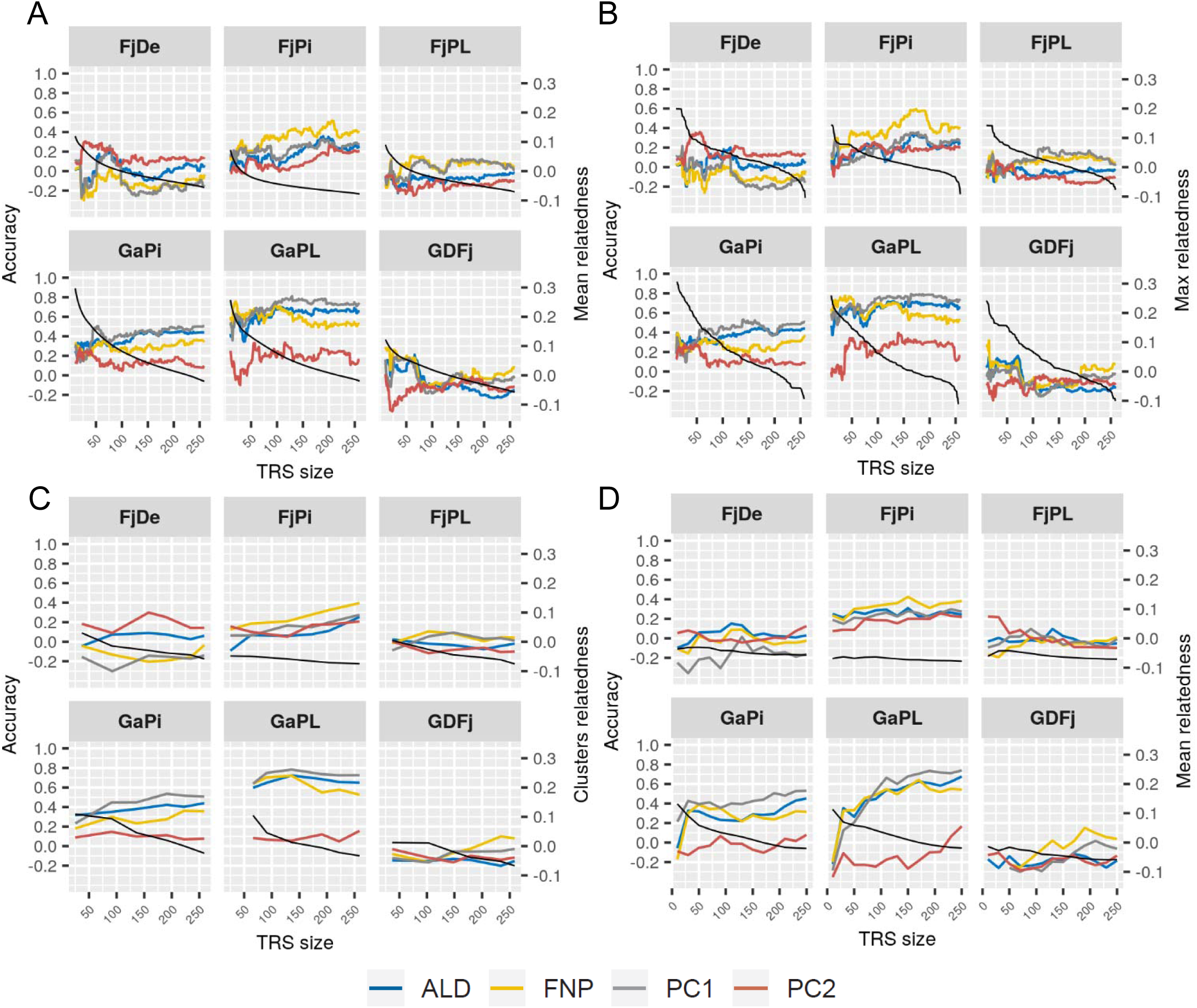
Optimization of the training population for the prediction of each family using *a priori* information on individuals. A, addition of individuals in the TRS by decreasing mean relatedness to the predicted family; B, addition of individuals in the TRS by decreasing maximum relatedness to the predicted family; C, addition of clusters by decreasing mean relatedness to the predicted family; D, selection of individuals for TRS of different sizes based on the five principal components obtained with discriminant analysis of principal components and using the CDmean design criteria. The color legend applies for all parts of the figure.

## Discussion

In this work we assessed the feasibility of genomic selection (GS) for apple texture by performing an in-depth analysis of this complex phenotype together with the genetic correlates influencing its genomic predictions. The results presented here on genomic prediction for apple texture evidenced a large potential for GS for this trait, providing important key elements and tools to set-up a prediction experiment given the available genetic information in any apple population.

### Family-dependent fruit texture profiles and fruit texture prediction

The texture dissected “sub-traits” were highly heritable, although variability within families was very contrasted, showing, in specific cases, a transgressive segregation, such as ‘FjPL’. Although the traits were predictable with moderate to high accuracy within the collection (accuracies between 0.41 and 0.64), this was not easily achievable in all biparental families. Without TRS optimization, texture could be accurately predicted for ‘GaPL’ (mean accuracy of 0.57), while ‘GaPi’ and in ‘FjPi’ showed a moderate prediction accuracy (mean accuracy of 0.30). In contrast, near-zero or negative accuracies were instead obtained for ‘FjDe’, ‘FjPL’ and ‘GDFj’ across traits (mean accuracy of −0.05). Surprisingly, large negative accuracy values were repeatedly obtained in ‘FjDe’ and ‘GDFj’, which could be potentially explained by the strong epistatic effect possibly present in these families (Lehner, 2011) or by a systematic bias due to the calculation of the Pearson correlation coefficient (Zhou *et al.*, 2016), indicating that fruit texture cannot be predicted in these families using the entire collection as TRS. In contrast, previous works on firmness and crispiness yielded mostly low accuracies when predicting unobserved genotypes in a set of families or in a collection (between 0.15 and 0.35, Kumar *et al.* 2015, McClure et al. 2018). A much higher accuracy of 0.83 was found for firmness by Kumar *et al.* (2012), which can be mainly explained by their crossing design and validation procedure. In the present study, the analysis of PCA allowed to better understand the relation between firmness and crispiness, both positively correlated and summarized by PC1 and PC2, with PC2 specifically dissecting the difference between these two texture sub-traits. When used as synthetic trait in the computation, PC1 was among the best predictable traits (accuracy of 0.59 in collection and highest accuracy among traits and family: 0.73 in GaPL), justified by the 80.5% of total phenotypic variation explained by PC1, while PC2 accounted only for 12.7%. Despite the lower variability of PC2, this trait could be predicted with a reasonable accuracy of 0.42 in the collection, while in most of the families the accuracy level was above 0.2 (with, and in some cases without TRS optimization). PC2 was not predictable in ‘GDFj’ and ‘FjPL’, two families with moderate and high transgression on the PC2 axis. The results showed that using PC1 and PC2 as a first tentative to perform a multi-trait prediction was a relevant method to predict fruit texture profiles through an integrative approach.

### Impact of genetic clustering and relatedness on prediction accuracy

Having highly related individuals between the TRS and the TS is necessary but not always sufficient for an optimal TRS design; in fact enlarging the TRS with scarcely related individuals can diminish prediction accuracies (Lorenz & Smith, 2015). Moreover, trait variation can be coupled with genetic structure. Several studies have for instance showed the impact of genetic structure on genomic prediction, demonstrating that taking genetic structure into account can improve GS efficiency (Guo *et al.*, 2014; Isidro *et al.*, 2015; Rio *et al.*, 2019). Although in apple the genetic structure is known to be weak, with substantial levels of admixture in apple cultivars (Urrestarazu *et al.*, 2016; Vanderzande *et al.*, 2017; Cornille *et al.*, 2019), it could still have a relevant effect on predictions, depending on the population composition and the trait under investigation. Significant genetic structure has been identified, for instance, between dessert and cider apples, which could potentially be correlated with fruit quality traits (Lassois *et al.*, 2016). Through the implementation of the DAPC method, six significant although lowly differentiated genetic clusters were obtained, with families belonging to one or two specific clusters, depending mostly to the assignment of their parental cultivars. While some degree of correlation was apparent between the genetic clustering of individuals and their phenotypic distribution (Fig. S3), the addition of the clustering effect into the prediction model almost systematically degraded the prediction accuracies. Moreover, the TRS optimization based on clustering was the lowest performing among the four methods tested. This could indicate that additive relationship alone already captured the genetic clustering present in our population. One important information given by the clustering patterns was that the ‘GaPL’ and ‘GaPi’ families, for which both parents were in the same genetic cluster or in the best represented cluster in the collection (Cluster 5), yielded the best predictions.

The genetic parameter having the largest impact on predictions was genetic relatedness, with the two families most related to the collection (‘GaPL’ and ‘GaPi’) yielding by far the highest accuracies compared to the remaining Fuji-related families. This observation finds consistency to the fact that genetic relationship is a fundamental parameter in genomic prediction (see *e.g.* Habier *et al.*, 2010; Clark *et al.*, 2012; Daetwyler *et al.*, 2014). The addition of closely-related individuals from the same family (scenario 2) or from a complete half-sib family (scenario 3) to the collection did not improve the prediction accuracy, except for ‘FjPi’, for which scenario 2 was the most accurate. This result might indicate that either the collection retains already ‘enough’ diversity to predict families, or that the excess of unrelated individuals in the collection cannot be corrected by adding related individuals. Thus, scenario 2 and 3 do not seem to effectively improve the TRS.

To this end, the gradual increase of the TRS size using *a priori* information of genetic parameters was used as an alternative optimization strategy. TRS optimization was tested in four different ways, based on *a priori* information on similarities between individuals. These were represented either by additive relationship or by genetically derived principal components coordinates (Fig. 5, Fig. S7, Table 3, Table S7). The results allowed in all cases to improve predictions tested beforehand with TRS scenarios 1 to 3 with a minimal increase of 0.2 and maximal increase of 0.4, reaching a maximum accuracy of 0.81 (‘GaPL’, PC1, Table 3). This means that the maximum accuracies were also never reached by employing the entire collection, especially for families with the lowest genetic relatedness to the TRS (*i.e.* to the collection here). The best prediction accuracy for fruit texture in apple was obtained with the implementation of 50 individuals in the TRS for families less related to the entire TRS and at least 100 accessions for families with a higher genetic relationship (or clustering within the major genetic cluster of the TRS, such as ‘GaPL’ and ‘GaPi’ here). These results are consistent with previous findings in barley from Lorenz and Smith (2015), that showed the detrimental effects of adding unrelated individuals to the TS into the TRS, partially contradicting the idea that having at least one related individual in the TRS is sufficient to increase accuracies (Daetwyler *et al.*, 2014).

Our results thus provided useful information for the TRS composition, illustrating the complex roles of structure and relatedness in shaping texture variability in apple.

### Towards a simplified assessment of fruit texture for genomic selection

The improvement of fruit texture is still limited by the time-consuming and expensive assessment needed for its dissection and the low variation observed in modern elite apple accessions due to the fixation of PG1 (Atkinson *et al.*, 2012; Di Guardo *et al.*, 2017). Thus, even though we demonstrate the feasibility of GS for apple texture, its application will be considered only if predictions are precise enough to perform the costly phenotyping of the TRS. The characterization of texture is a challenging task, as this trait is composed of mechanical and acoustic sub-traits. The analysis of PC1 and PC2 relied on the texture dissection and the measurements of these 12 traits. In particular, FNP, which is the number of mechanical peaks observed in the mechanical profile generated by fruit compression on the texture analyzer, was highly correlated with the group of the acoustic traits related to crispiness. As mechanical traits are easier to measure than acoustic ones, FNP would be in practice the best measurement to choose for assessing crispiness. Since we also obtained high prediction accuracy for FNP (0.63 in collection and maximum of 0.78 in “optimized” family prediction), we propose this sub-trait as the most valuable descriptor for fruit texture, minimizing the effort needed to phenotype such as complex phenotype. Moreover, the predictions presented in this study have been performed with a set of 8,294 SNPs, which is still not dense enough considering the rapid decay of the linkage disequilibrium in apple (Laurens *et al.* 2018). Although we reached already satisfying accuracies with this amount of SNP, it would be useful to increase the number of markers with the available apple 480K (Bianco *et al.*, 2016) or by using genotyping-by-sequencing methods (Gardner *et al.*, 2014) to further improve predictions.

The use of principal component as synthetic traits resulted to be a valuable multi-trait approach to better predict and understand the texture variability. Here, we investigated in details that fruit crispiness (PC2) in particular is less variable than fruit firmness (PC1). While crispy apples are necessarily firm, the opposite relationship is in fact not validated. Our predictions indicated that fruit firmness in apple can be accurately selected (along PC1), but it needs to be taken into account that an excessive value for this trait can lead to unpleasant quality perception for the consumer. On the other side, crispiness was better predicted with the PC2. Despite the lower variation for crispiness in our population, the selection for this trait resulted to be feasible, although with lower accuracy. To improve the predictions for crispiness we might need to increase the variability for this trait within the TRS. More generally, while the selection on fruit traits has shaped apple domestication, the current cultivated pool relies on a few founders, hence having a narrow genetic basis. Thus, a better targeting of apple texture might necessitate a pre-breeding step incorporating or generating genetic diversity for this trait with the use of mealy cultivars and of wild relatives of *Malus domestica* (Khan *et al.*, 2014; Peace *et al.*, 2019).

## Supplementary data

**Table S1**. Texture genotypic values and coordinates for PC1 and PC2.

**Table S2**. Additive relationship matrix.

**Table S3**. Assignments of individuals to genetic clusters.

**Table S4**. Pairwise Fst-values between genetic clusters.

**Table S5**. Accuracies obtained in cross-validations within the collection using two models.

**Table S6**. Accuracies obtained in family predictions using two models and three TRS scenarios.

**Table S7**. Accuracies obtained in family predictions with TRS optimization with four methods.

**Fig. S1**. Bayesian information criterion values obtained in the discriminant analysis of principal components.

**Fig. S2**. Distribution of texture genotypic values according to the type of population.

**Fig. S3**. Distribution of texture genotypic values according to cluster assignments.

**Fig. S4**. Accuracies obtained from cross-validations within the collection.

**Fig. S5**. Accuracies obtained for each family in three prediction scenarios with model B.

**Fig. S6**. Observed vs. predicted values in predictions for each family on each trait with model A.

**Fig. S7**. Comparison of methods for the optimization of the training population.

## Abbreviations

ALD: Acoustic Linear Distance
APMax: Acoustic Max Pressure
APMean: Acoustic Mean Pressure
ANP: Number of Acoustic Peaks
BIC: Bayesian Information Criterion
BLUP: Best Linear Unbiased Predictor
COLL: Collection
DAPC: Discriminant Analysis of Principal Components
FF: Final Force
FLD: Force Linear Distance
FI: Initial Force
FMax: Max Force
FMean: Mean Force
SNP: Single Nucleotide Polymorphism
PC: Principal Component
PCA: Principal Component Analysis
FNP: Number of Force Peaks
TS: Test Set
TRS: Training Set
YM: Young Module

## Acknowledgements

This work was co-funded by the EU seventh Framework Programme by the FruitBreedomics Project No. 265582: Integrated Approach for increasing breeding efficiency in fruit tree crops (www.FruitBreedomics.com). The views expressed in this work are the sole responsibility of the authors and do not necessarily reflect the views of the European Commission.

